# Detection of *Fusarium Oxysporum* F. SP. Elaeidis Causing Fusarium Wilt of Oil Palm Using Loop-Mediated Isothermal Amplification (Lamp)

**DOI:** 10.1101/540393

**Authors:** Kwasi Adusei-Fosu, Matthew Dickinson

## Abstract

*Fusarium oxysporum* f. sp. *elaeidis* (FOE) a pathogen that causes fusarium wilt disease in oil palm can be detected using polymerase chain reaction (PCR) but very time consuming. Loop-Mediated Isothermal Amplification (LAMP) was used to rapidly detect *Fusarium oxysporum* f. sp. *elaeidis* (FOE) in oil palm seedlings. Eight additional *Fusarium oxysporum* isolates collected from symptomatic oil palm trees (i.e. presumed-FOE as their pathogenicity was not confirmed) and five other non-FOE isolates were sampled from symptomatic mature oil palm trees and tomato respectively to broaden the scope of the research. The identities of FOE, presumed-FOE and non-FOE were established via sequencing. LAMP primers designed for detecting FOE or presumed-FOE were based on partial sequences of *Secreted In Xylem* (*SIX8*) and *P-450* cytochrome. The earliest detection time for *SIX8* and *P-450* cytochrome primers were 4:00 mins and 6:45 mins respectively with both recording late time for detection at 26:30 mins. Annealing derivative curves were used for assessing the level of specificity for both *SIX8* and *P-450* cytochrome, but none of the LAMP primers could distinguish between FOE, presumed-FOE and non-FOE.

## Introduction

The use of the LAMP could aid on-site detection of diseased plants [1] which drastically reduces the quantity of samples to be transported to the laboratory. This marked the beginning of several attempts made to develop real-time PCR equipment for use in the field [12, 22, 26].

LAMP is a rapid amplification method employing a strand displacing *Bst* DNA polymerase and 4 – 6 primers, two of which are ‘fold back’ primers [14, 19] which form stem-loop motifs with self-priming capability (Fig. 1.0). The primers used are two sets, the internal primers and external primers. Subsequent studies have found the use of additional ‘loop primers’, which bind to the loop structures and greatly reduce the reaction times [14], resulting in a total of 6 primers. The 60 – 65 °C reaction temperature combined with a minimum of four primers makes LAMP a highly specific reaction. The high level of specificity results from the requirement for primers to bind up to eight regions of the target sequence. This results in an amplification scheme where the priming sequence is copied with each round of replication and remains tethered to the previous amplicon resulting in a concatenated product of alternating sense / anti-sense repeats of varied length. This results in large amounts of amplicons which can be used for further studies in detection [5].

**Figure. 1.**
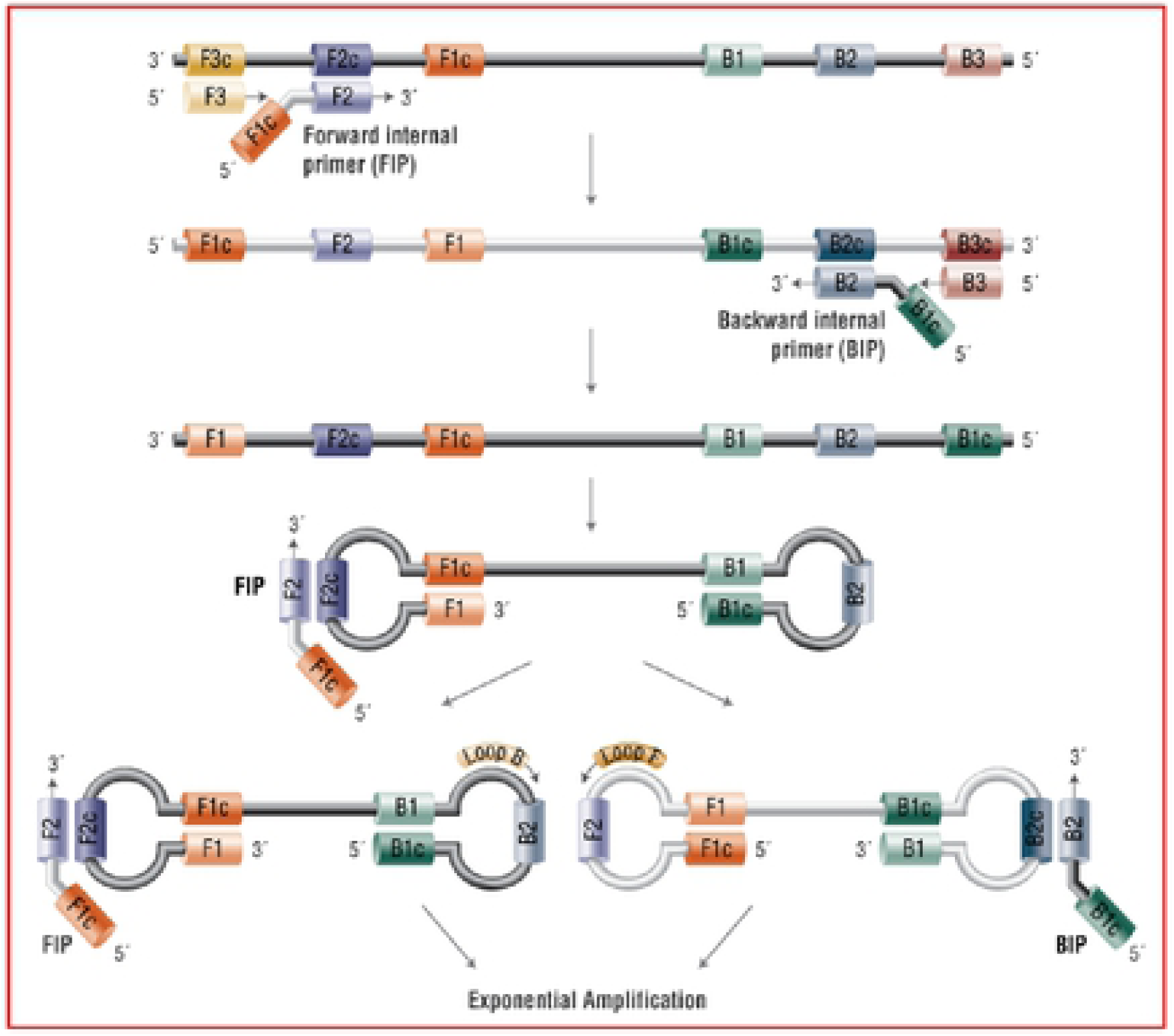
Loop-mediated Isothcnnal Amplification (LAMP) uses 4 - 6 primers recognizing 6 - 8 distinct regions of target DNA. A strand-displacing DNA polymerase initiates synthesis and 2 of the primers form loop structures to facilitate subsequent rounds of amplification (Source:https://www.neb.com/applications/dna­amplification-and-pcr/isothcnnal-amplification).

LAMP is one of the most well established methods for isothermal amplification of nucleic acids to date. The technique has been used as a molecular tool for the detection of several plant pathogens over recent years [7, 16, 26] including fungi [11, 18, 24]. There are several reports on LAMP for detecting *Fusarium* spp. [4, 6, 8, 17, 18]. LAMP assay could detect and differentiate *F. oxysporum* f. sp. *lycopersici* (*Fol*) race 1 isolates based on the *SIX4* and *SIX5* genes using three primer sets [4]. The usefulness of the analysis of fungal cultures by direct analysis of surface scrapings from agar plate cultures, direct testing of single infected barley grains, and detection of *Fg* in total genomic DNA isolated from bulk samples of ground wheat grains has been demonstrated [18]. LAMP has been used to successfully quantify genomic DNA of *F. oxysporum* f. sp. *cubense* (*Foc-*TR4) in soil samples [29]. The sensitivity of the LAMP has also been reported [3]. Even though PCR and LAMP assays would successfully detect positive infected samples of tomato with *Fol*, considering the time, safety, cost and simplicity, the latter technique was overall superior [4]. In addition, the real-time application with the Optigene-system (Optigene, UK) has several advantages such as easy mobility or portability of the detection device. This makes it portable for field work. Unfortunately no report has been published on LAMP tool/assay to detect isolates of *Fusarium oxysporum* f. sp. *elaeidis* (FOE). Hence, there was the need to develop LAMP primers to detect FOE in inoculated oil palm seedlings and *Fusarium oxysporum* isolates collected from symptomatic mature oil palm trees (i.e. presumed-FOE as their pathogenicity had not been tested although they were collected from symptomatic oil palm in the field) to enable faster screening of oil palm seedlings prior to transplanting them to field to ensure disease free oil palm seedlings were planted.

## Materials and methods

### DNA extraction, PCR and DNA sequencing

A total of 40 strains were used including eight *Fusarium oxysporum* isolates (presumed-FOE) collected from symptomatic oil palm mature trees (*Elaeis guineensis*), four FOE (BOP-B5, NORP-N5, OPRI-5 and 16F) isolates confirmed to be pathogenic against oil palm seedlings and the remaining non-FOE isolates collected from tomato. The identities of all the isolates were confirmed via sequencing. Mycelia (50 - 100 mg) from cultured FOE, presumed-FOE and non-FOE were isolated from PDA plates using a sterile surgical blade. Tissue disruption was carried out using glass beads and homogenizer (FastPrep®) at a speed of 6.5 ms^-1^ for a total time of 45 s in the presence of liquid nitrogen. DNA extractions of the cultures were then carried out using DNeasy Plant Mini Kit (Qiagen) according to the manufacturer’s protocol. Polymerase chain reaction (PCR) of various regions of the template DNA was performed using primer pairs of interest (Table. S1). PCR was carried out in 30 µl volumes consisting of 15 µl of master mix (MangoTaq™ DNA Polymerase), 1 μl (of 10 pmol / ul) each of all primer pairs mentioned in separate reaction mixtures, 12 µl sterile distilled water and 1 µl of template DNA of the isolates of interest. The reaction was performed in a BIO-RAD S1000 Thermal Cycler with the amplification conditions of 95 °C for 2 min for initial denaturation, followed by 35 cycles of denaturation at 95 °C for 2 min, annealing at temperatures suitable for amplification for each primer pair of interest and extension/elongation at 72 °C for 1 min 30 sec. The final extension was set at 72 °C for 5 min. PCR products were cleaned using the QIAquick PCR Cleanup kit (Qiagen) following manufacturer’s instruction followed gel electrophoresis using 1 kb ladder (Promega). Sequencing reactions were performed by Fisher Scientific or MyGATC. BLAST searches were performed using the GenBank sequence database to confirm the identity of the fungal isolates sequences based on the *Secreted In Xylem* (*SIX*) gene and *P450 cytochrome oxidase* used for amplifying the rDNA. The output from BLAST algorithms was used to query any unknown sequences against the database of all the fungal gene regions. These sequences were subsequently used to design LAMP primers.

### Loop-Mediated Isothermal Amplification (LAMP) primer design

#### Loop Mediated Isothermal Amplification (LAMP) assay

The LAMP primers (Table S1) were self-designed from partial sequences based *Secreted In Xylem* (*SIX8*, *SIX10* and *SIX13*) gene and *P450 cytochrome oxidase. Fusarium oxysporum* f. sp. *elaeidis* (FOE) detection assay was done by preparing the LAMP primer mix which consisted of 152 µl sterile distilled water with primer concentration of 10 µM each of the F1, B1, F2, B2 primers and 2 µM each of FIP, BIP primers. Master mix for eight reactions was prepared which consisted of 23 µl of LAMP primer mix, 46 µl sterile distilled water and 115 µl Optigene master mix (dNTPs, *Bst* DNA polymerase and MgCl_2_). A final volume of 21 µl in each of the eight LAMP tubes consisted of 20 µl of the reaction mix dispensed into each of the LAMP tubes and 1 µl genomic DNA. The detection time was set to 30 min for all reactions for each primer set.

### Results for Loop-Mediated Isothermal Amplification (LAMP)

LAMP assays were developed for two sets of genes, those encoding *Secreted In Xylem* (*SIX*) and the cytochrome *P450*. Results to confirm the presence or absence of FOE as well as detection time in genomic DNA of all isolates used for the study are shown in Tables 2 & 3. The amplification and derivative curves generated were also observed to confirm the specificity of the products amplified (example is shown in Fig. 2). Generally, amplification was observed at 65 °C. *SIX8* gene could amplify all FOE, some presumed-FOE and non-FOE isolates (Table 2). *SIX10* and *SIX13* could not detect FOE and presumed-FOE isolates. Generally, the detection times varied among all the genes (*SIX8* or *P450*) that were used (Tables 2 & 3). The time of detection for all the *SIX* genes was between 4:00 min to 29:15 min. *P450 cytochrome* detected isolates at 6:45 min (Table 3). Both *SIX8* and P450 *cytochrome* LAMP primers designed recorded a detection time below 30 min in FOE and presumed-FOE isolates. The LAMP primers were randomly tested on the sampled roots of oil palm seedlings that showed symptoms of FOE infection to confirm the sensitivity of the assay for on-site detection, and these assays were positive with the *SIX8* gene primer set but not consistent, hence results not shown.

## Discussions

The research showed that LAMP primers could detect both FOE within inoculated oil palm seedlings and presumed-FOE (symptomatic oil palm in fields). The time for detection FOE or presumed-FOE using *SIX* gene and P450 *cytochrome* primers differed but all could detect either FOE or presumed-FOE within 30 min compared to PCR which is time consuming. Although there are reports on Loop-Mediated Isothermal Amplification (LAMP) assays for detecting several *formae speciales* (f.spp.) for *Fusarium* [1, 4, 8] and other plant pathogens [27] this is the first time an attempt has been made to use LAMP for detection of *Fusarium oxysporum* f. sp. *elaeidis* (FOE).

*Secreted In Xylem (SIX*) gene, are small, secreted and well known to be cysteine-rich, first identified in the xylem sap of tomato infected with *Fol* [9, 23]. Loop Mediated Isothermal Amplification (LAMP) was successfully designed based on *SIX* genes to distinguish *Foc* from other plant pathogenic fungi [4, 27]. The detection of *Fol* with LAMP was achieved based on the 28S rRNA regions [3, 27] but unable to distinguish between pathogenic races of *Fol* isolates. The LAMP assay using the *SIX8* was positive for FOE, presumed-FOE and non-FOE. The presence or absence of some of the *SIX* genes in FOE, presumed-FOE and non-FOE used in this study is congruent with a study that showed that as of now, only fourteen *SIX* (1-14) genes have been identified and most share similarities with each other or with other fungi [28]. The LAMP assay for *SIX8* was faster for detection but detection time varied from one isolate to the other and differed as well among FOE, presumed-FOE and non-FOE. This could be because of the differences in the genomic DNA concentrations used in the study and the presence of some inhibitors as well which influenced the time of amplification. LAMP *SIX8* primer in this study, could not directly detect FOE (OPRI-5, BOPP-5, NORP-5 and 16F) isolates that were artificially inoculated into soil. On the contrary, other research could directly detect *Fol* race 1 in soil artificially inoculated based on primers designed for *SIX4* and *SIX5* [4]. Similarly, LAMP as an effective tool for detecting *Foc* race 4 isolate in soils has been reported [20].

It is reported that the *P450 cytochrome* are distributed widely in many organisms [15]. *P450 cytochrome* has been associated with pathogenicity in some fusarium such as the *F. oxysporum* f. sp. *cubense* (*Foc*) [25]. In this study, *P450 cytochrome* was detected in FOE, presumed-FOE and non-FOE isolates. The level of *P450 cytochrome* differences such as the copy numbers or gene families, significantly varies biologically across kingdoms, phyla and species [15]. Furthermore, *P450 cytochrome* share conserved overall protein architecture and have many conserved sequences, despite the higher level of diversity in the *P450 cytochrome* [15]. These characteristics of the *P450 cytochrome* may have contributed to the presence in FOE, presumed-FOE and non-FOE as well as the varying time of detection. The variation in time of detection using *P450 cytochrome* LAMP primers could be because the differences in the genomic DNA concentrations isolated from FOE, presumed-FOE and non-FOE.

Generally, LAMP primers developed in this study, either the *P450 cytochrome* or *SIX8* genes represents an extremely rapid [10] diagnostic tool for FOE. However, as at now, the assays lack the specificity required to discriminate between FOE, presumed-FOE and non-FOE isolates. However, the primers provided could potentially be used to detect or screen FOE as it is host specific to oil palm seedlings especially in nurseries to prevent the spread and introduction of the pathogen in various oil palm plantation sites.

## Acknowledgement

This research was fully funded through Commonwealth Scholarship Commission-United Kingdom and I sincerely thank them for the immense excellent and outstanding support. We also thank Council for Scientific and Industrial Research – Oil Palm Research Institute, Ghana for supplying oil palm seedlings for the study. We honestly thank Dr. Julie Flood the Global Director of CABI Research for technical advice and support and the late Professor Richard Cooper former Lecturer at University of Bath – United Kingdom for providing us with two isolates of *Fusarium oxysporum* f. sp. *elaeidis*. Further thanks go to Dr. Y. Ndede a Scientist and Frank Dwumfour a Technician at the Council for Scientific and Industrial Research - Oil Palm Research Institute, Ghana, for the immense assistance during the field work and sampling. Our appreciation also goes to the School of Biosciences, Plant and Crop Science Division, The University of Nottingham – United Kingdom, for giving us all laboratory resources needed for this research. Our final appreciation goes to the various oil palm plantation sites where sampling was undertaken including NORPALM-Ghana, BOPP-Ghana and TOPP-Ghana.

---

**Table 1:** Lisi of selected *Fusarium orysporum* isolates collected from symptomatic oil palm and other fusarium isolates (non-FOE)

| *Fusarium* species | Isolate code | Location |
| --- | --- | --- |
| **\**F.oxysporum* f. sp. *elaeidis*** | IVORY COAST-16F | Ivory Coast |
| **\**F.oxysporum* f. sp. *elaeidis*** | DR CONGO-F3 | DR Congo |
| *F.oxysporum* f. sp. *elaeidis* | DR CONGO-Z/CBS217.49 | DR Congo |
| *F.oxysporum* f. sp. *elaeidis* | SURINAME-S/CBS783.83 | Suriname |
| *F.oxysporum* f. sp. *elaeidis* | GHANA NORPALM-N2 | Ghana |
| *F.oxysporum* f. sp. *elaeidis* | GHANA NORPALM-N3 | Ghana |
| **\**F.oxysporum* f. sp. *elaeidis*** | GHANA NORPALM-N5 | Ghana |
| *F.oxysporum* f. sp. *elaeidis* | GHANA BOPP-B4 | Ghana |
| *F.oxysporum* f. sp. *elaeidis* | GHANA BOPP-B5 | Ghana |
| **\**F.oxysporum* f. sp. *elaeidis*** | GHANA OPRI-5 | Ghana |
| *F.oxysporum* f. sp. *elaeidis* | GHANA OPRI-11 | Ghana |
| *F.oxysporum* f. sp. *elaeidis* | GHANA OPRI-18 | Ghana |
| *F.oxysporum* sp. | Fus126B-3 | ADAS |
| *F.oxysporum* sp. | Foxy150-5 | ADAS |
| *F.oxysporum* sp. | Foxy sp-8 | ADAS |
| *F.oxysporum* sp. | Foxy2571-9 | ADAS |
| *F.oxysporum* sp. | Foxy sp-10 | ADAS |
| *Fusarium* sp. | Fus-11 | ADAS |
| *F.oxysporum* sp. | FoxyBX14/153b-A | ADAS |
| *F.oxysporum* sp. | FoxyBX14/153a-B | ADAS |
| *F.oxysporum* f. sp. *lycopersici* | Foxy.Forla-C | UoN |
| *F.oxysporum* f. sp. *lycopersici* | Foxy.Forlb-D | UoN |
| *Fusarium solani* 1 | F1 | UoN |
| *Fusarium solani* 2 | F2 | UoN |
| *Fusarium oxysporum* f.sp. *radicis-lycopersici* | F3/ FORL A | UoN |
| *Fusarium oxysporum* f.sp. *radicis-lycopersici* | F4/ FORL B | UoN |
| *F.oxysporum* BX15/4a | F5 | UoN |
| *F.oxysporum* BX15/4b | F6 | UoN |
| *F.oxysporum* BX15/4c | F7 | UoN |
| *F.oxysporum* f. sp. *cubense* (FOC2A) | F8 | UoN |
| *F.oxysporum* f. sp. *cubense* (FOC2B) | F9 | UoN |
| *F.oxysporum* BX14/153a | F10 | UoN |
| *F.oxysporum* BX14/153b | F11 | UoN |
| *F.oxysporum* BX14/153c | F12 | UoN |
| *F.oxysporum* BX14/152a | F13 | UoN |
| *F.oxysporum* BX14/128a | F14 | UoN |
| *F.oxysporum* BX14/168d | F17 | UoN |
| *F.oxysporum* f. sp. *lycopersici* race 1 | F18 | Alison Jackson (UK) |
| *F.oxysporum* f. sp. *lycopersici* race 2 | F19 | Alison Jackson (UK) |
| *F.oxysporum* f. sp. *radices lycopersici* | F20 | Alison Jackson (UK) |

**\**F.oxysporum* f. sp. *elaeidis*** = FOE (i.e. Pathogenic to oil palm)
*F.oxysporum* f. sp. *elaeidis* = presumed-FOE (i.e. Pathogenicity not confirmed in oil palm though sampled from symptomatic oil palm trees)

